# A bioinformatic workflow to facilitate the study of less-understood proteins: the case of SH2D2A

**DOI:** 10.1101/2025.02.11.637442

**Authors:** Brian C. Gilmour, Hanna Chan, Sachin Singh, Tuula A. Nyman, Andreas Lossius, Paweł Borowicz, Santosh Phuyal, Anne Spurkland

## Abstract

Adaptor proteins are key regulators of immune signalling but are challenging to study due to redundant pathways that obscure clear functional “anchor” phenotypes. Making use of public single-cell RNA sequencing (scRNA-seq) datasets, we theorised that anchors could be identified by working in the reverse direction: starting from the transcriptomic level and working “bottom-up” to identify anchoring phenomena at the cell surface. We have produced a prototype workflow for this method using the case of the lymphocyte-enriched adaptor protein SH2D2A, whose function remains uncertain. Using our bottom-up analysis, we have identified a link between SH2D2A and T cell-T cell (T-T) synapses enriched for ITGB2. The functional implications of this link are currently being explored, but further application of this bioinformatic approach to less-understood proteins may prove useful in understanding the minutiae of immune cell signalling, with implications for immunotherapy.

---

Lymphocytes play key roles in immune defence, relying on both intrinsic and extrinsic signals to coordinate responses. While much progress has been made in understanding cytokine^**1–4**^ and pathogen-recognition receptor signalling^**5–10**^, as well as pathways downstream of specific receptors^**11–14**^, many lymphocyte signalling mechanisms remain incompletely characterized. Advances in these areas have already led to novel immunotherapies, particularly for tumour eradication. However, numerous unexplored pathways and molecules could be leveraged for therapeutic purposes if their downstream effects were better understood.

Adaptor proteins are an underexplored but essential family, particularly in effector lymphocytes, where they integrate immune signals to guide responses^**15,16**^. However, studying them is challenging since they often participate in multiple pathways, with their loss masked by compensatory mechanisms^**17,18**^. Traditional research relies on “anchors”—clear, measurable phenomena—to trace molecular functions, working top-down from observable effects to proteins and transcripts. This approach has revealed key roles of important proteins, such as LAT’s function as a focal point in T cell receptor (TCR) signalling^**19,20**^.

While this conventional top-down approach remains a gold standard in research, it is less-suited for studying adaptor proteins due often to their subtle phenotypes and lack of clear anchors. Instead, a “bottom-up” approach—starting with transcriptomic data, identifying co-expressed proteins, and inferring potential functional anchors—may be more effective. The growing availability of high-quality single-cell RNA sequencing (scRNA-seq) datasets has made this method increasingly feasible.

To test this approach, we developed a bioinformatic workflow using SH2D2A, an adaptor protein enriched in T and natural killer (NK) cells, as a case study. SH2D2A is upregulated upon activation^**21–23**^ and is also expressed in some epithelial cells^**24–26**^. It interacts with several kinases^**24,25,27–32**^, including the kinases ITK and LCK in immune cells, and SRC in epithelial cells.

In epithelial cells, SH2D2A modulates VEGF signaling *via* SRC^**24–26**^, but its role in immune cells remains unclear. Though linked to TCR signaling through LCK^**23,31**^,SH2D2A knockout studies show minimal effects on T cell function^**31–33**^, leaving its precise role uncertain. This lack of clear anchors makes SH2D2A an ideal candidate for our anchor-generating bioinformatic analysis. In applying this bottom-up strategy, we aimed to identify functional anchors around which to base further study of SH2D2A.

Applying our bottom-up approach, we identified SH2D2A as being connected to cytotoxicity and integrin-mediated cell-cell adhesion. Translating these findings into the wet-lab, we found integrin-family proteins enriched in SH2D2A pulldowns from primary human T cells *via* mass spectrometry. Additionally, we found that SH2D2A localizes to T-T junctions defined by ITGB2 alongside LCK but not CD3ζ, suggesting it plays a role in TCR-independent signaling at integrin-associated junctions. This suggests that our workflow is capable of deriving novel information and new anchor proteins and phenomena for understudied gene/protein pairs working bottom-up from transcriptomic data.

## Results

### NK cells, memory CD8+ T cells, and innate-like T cells are the main expressors of SH2D2A in the blood

The role of SH2D2A in immune cells remains unclear, despite its known interaction with LCK, a key component of TCR signalling. Using proximity ligation assay (PLA), we confirmed the interaction of SH2D2A and LCK in primary T cells but found no stimulation-dependent changes in interaction, in contrast to PLA results for CD3ζ-ZAP70 (**Figure S1**). This finding obfuscated further research into SH2D2A’s role in TCR signalling and, lacking other credible candidates to anchor further research on, we turned to using SH2D2A as a test protein in a bioinformatic workflow that could attempt to predict novel anchors and areas of function bottom-up from transcriptomic data.

We selected the Liu *et al*. (*Cell*, 2021)^**34**^ dataset, containing adaptive (247K cells) and innate (125K cells) immune cells, and focused on T and NK cells. These were isolated and combined into a modified dataset (**Figure 1A-B**). To infer surface protein expression, we integrated this dataset with ADT data from Hao *et al*. (*Cell*, 2021)^**35**^ using *Seurat*, creating the mapped (MAP-Liu) dataset (**Figure 1C**).

**Figure 1.**
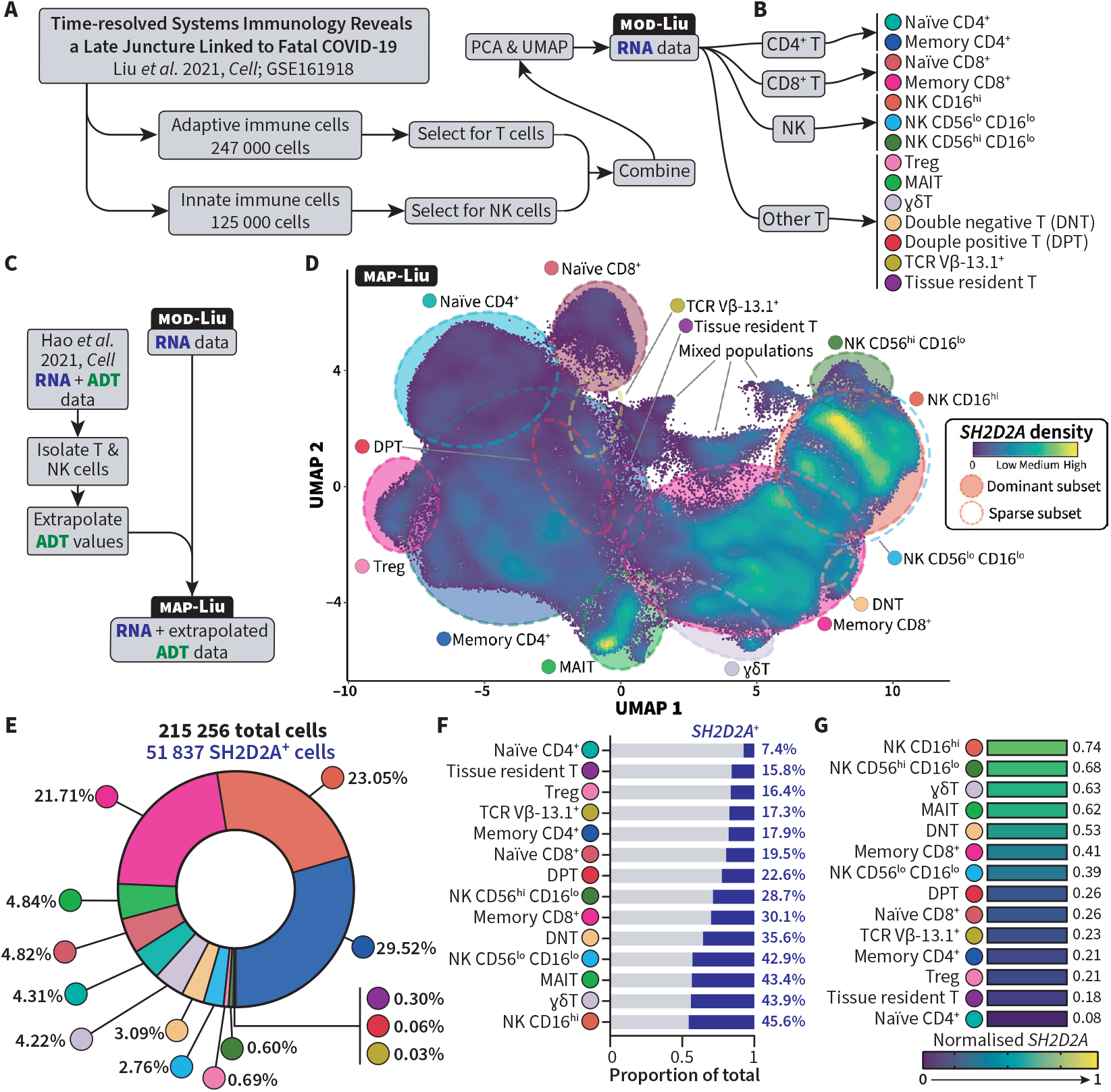
NK cells, memory CD8^+^ T cells, and innate-like T cells are the main expressors of *SH2D2A* in the blood. **(A)** Workflow to produce the modified Liu *et al*. dataset from the original data (GSE161918), consisting of only T and NK cells, and **(B)** the two layers of cell annotation metadata available in the modified Liu *et al*. dataset. **(C)**Workflow to map the ADT assay from the PBMC reference dataset Hao *et al*. onto the modified Liu *et al*. dataset to produce the mapped Liu *et al*. dataset, consisting of its original RNA assay and extrapolated ADT assay data. **(D)** UMAP of the mapped Liu *et al*. dataset (MAP-Liu), showing the density of the expression of *SH2D2A* and annotated with the general locations of each layer 2 cell type. **(E)** Summary of the proportion of the total SH2D2A^+^ cells represented by each cell type. Cell types are order counterclockwise from largest SH2D2A^+^ total. (F) The proportion of SH2D2A^+^ positive cells within each cell type, ordered from smallest (top) to largest (bottom), the average for the whole dataset is 24.1%. **(G)** Average *SH2D2A* expression in each cell type, ordered from highest (top) to lowest (bottom), the average for the dataset is 0.38. Expression levels were normalised to the highest expression value in the dataset.

SH2D2A was predominantly expressed in NK, memory CD8^+^, and innate-like T cells (**Figure 1D**), with highest proportions and expression levels in these subsets (**Figure 1E-G**), in contrast to CD4^+^ T cells, which had previously served as the main model of study.

### A sub-population of naïve CD8+ T cells express SH2D2A and are transcriptionally distinct from naïve SH2D2A– CD8+ T cells

To determine if SH2D2A marked any specific cellular states, we performed differential gene expression analysis between SH2D2A^+^ and SH2D2A^−^ populations within each cell type, with the results visualised as a volcano plot (**Figure 2A**). While NK, innate-like T, and memory CD8^+^ T cells showed minimal differences, naïve CD8^+^ T cells exhibited abundant differentially expressed genes and proteins contrasting the SH2D2A^+^ and SH2D2A^−^ populations (**Figure 2B**).

**Figure 2.**
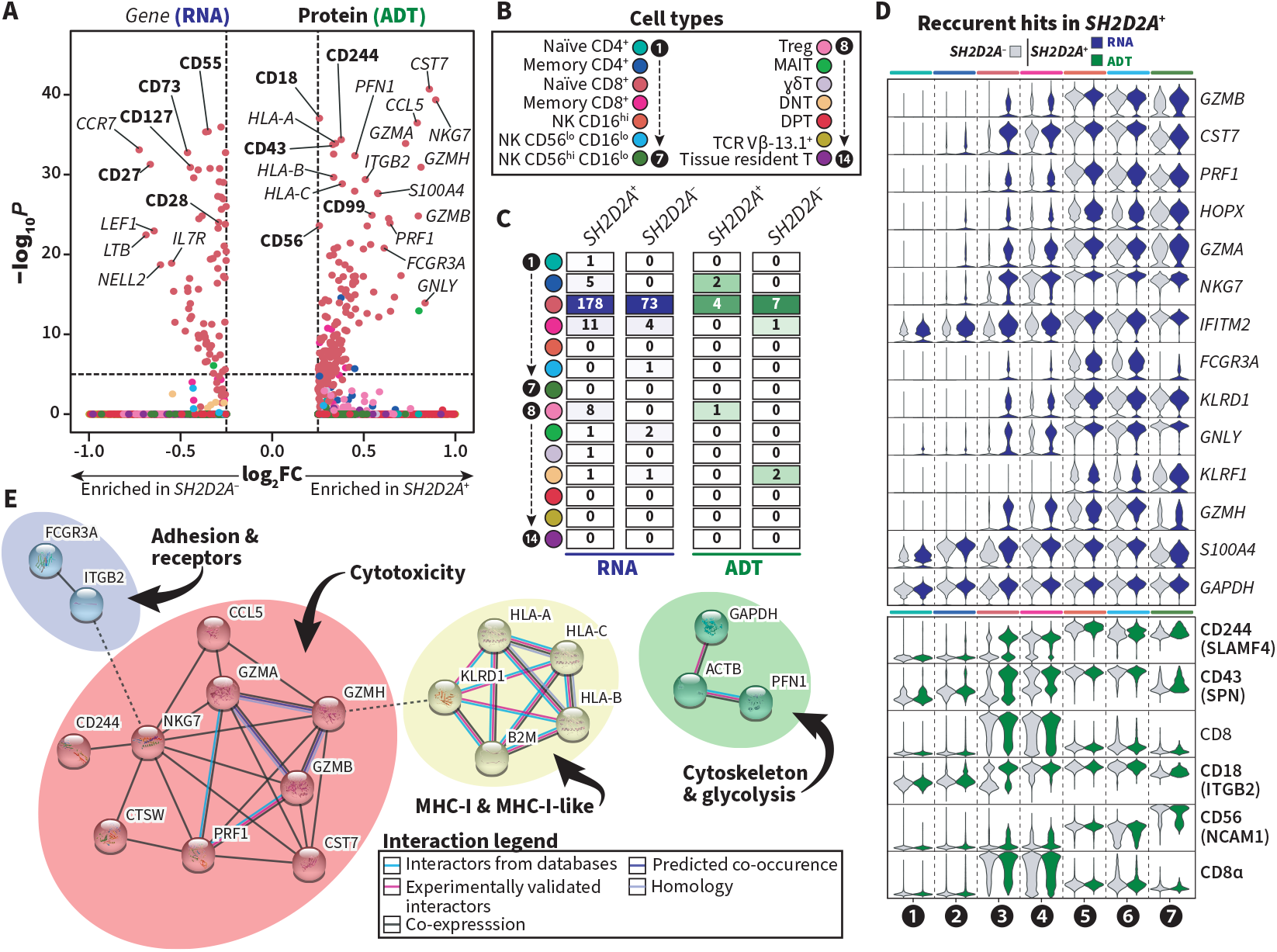
A sub-population of naïve CD8^+^ T cells express SH2D2A and are transcriptionally distinct from naïve SH2D2A^−^ CD8^+^ T cells. **(A)** Volcano plot of differential expression analysis of the *SH2D2A*^+^ vs. the *SH2D2A*^−^ portions of each cell type, both differentially expressed proteins **(bold)** and genes (*italics*) are shown, Wilcoxon ranked-sign test with Bonferroni correction was used to determine statistical significance. (B) colour legend for cell types used in **(A), (C)**, and **(D). (C)** Total numbers of differentially expressed genes (blue) and proteins (green) expressed in the SH2D2A^+^ and SH2D2A^−^ portions of each cell type. **(D)** Violin plots of differentially expressed genes (blue, upper) and proteins (green, lower) that are upregulated in the *SH2D2A*^+^ portion of >1 cell type. **(E)** STRING network following MCL clustering of the top 25 most significantly upregulated genes and proteins in naïve *SH2D2A*^+^ CD8^+^ T cells, unconnected nodes have been hidden, MCL clusters have been manually annotated based on known and common functions of their constituent elements.

To get a better picture of which genes re-occurred in several analyses, we summarised the genes (RNA, blue) and proteins (ADT, green) significantly upregulated in SH2D2A^+^ cells across multiple cell types (**Figure 2C**), the results included a cohort of cytotoxicity-related genes (*GZMB, CST7, PRF1, etc*.) and cytotoxicity-linked proteins (CD244, CD43, CD18, CD8). STRING^**36**^ network analysis of the top 25 differentially expressed elements in naïve CD8^+^ T cells, clustered using MCL^**37**^, highlighted cytotoxicity-associated transcripts and receptors like CD16 (*FCGR3A*) and CD18 (*ITGB2*) (**Figure 2D**).

### SH2D2A expression correlates with markers of cytotoxicity and cell-cell adhesion

To further explore elements co-expressed with SH2D2A, we increased clustering resolution to 27 clusters (**Figure 3A**) and analyzed their overlap with annotated cell types (**Figure 3B**).

**Figure 3.**
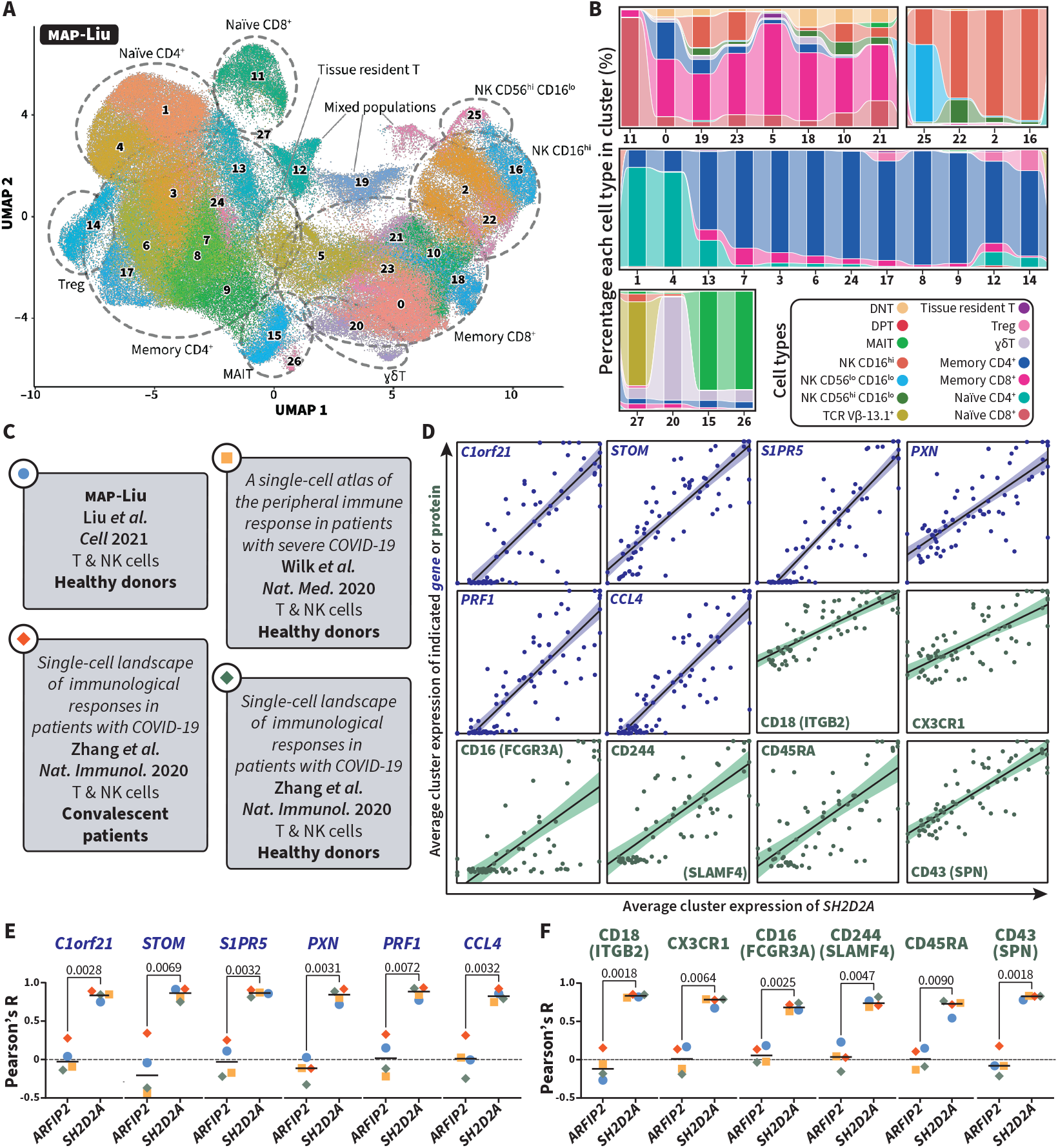
Expression of *SH2D2A* is correlated with expression of markers of cytotoxicity and cell-cell adhesion across four datasets. **(A)** UMAP of the mapped Liu *et al*. dataset following unsupervised clustering at higher resolution, showing all 27 produced clusters, the UMAP has been annotated to show the primary locations of the different cell types. **(B)** Alluvial plot of the proportions of each annotated cell type in each of the 27 cell clusters, clusters are split into 4 groups as produced by cutting the hierarchical clustering tree at its most distant branches (see **Figure S2**). **(C)** Summary of the 4 datasets used in cluster correlation analysis. **(D-E)** The top 6 genes **(D)** and proteins **(E)** with high correlation to *SH2D2A* expression, each dot represents a discrete cluster from one of the 4 datasets, clusters from all datasets were pooled for initial correlation analysis. **(F-G)** Consensus analysis of the top 6 most-correlated genes **(F)** and proteins **(G)** identified in each of the datasets in **(C)**, each dot represents the Pearson’s R from the correlation analysis of its own clusters. A paired, parametric, two-tailed Student’s *t*-test was used to determine the statistical significance of each gene or protein’s correlation to *SH2D2A* vs. a gene selected by random number generation (*ARFIP2*).

To minimize dataset-specific bias, we incorporated three additional datasets: healthy donors from Wilk *et al*. and Zhang *et al*., as well as convalescent patients from Zhang *et al*. (**Figure 3C**).We applied the same filtering and mapping procedures as in the MAP-Liu dataset (**Figure 1A & C**).

Averaging expression of all genes and mapped proteins across each cluster, we conducted correlation analysis to identify genes and proteins with a similar expression pattern to SH2D2A. The top six correlated genes (blue) and proteins (green, **Figure 3D**) included cytotoxicity-related markers (CD244, *PRF1*) alongside migration (*CCL4*, CXCR1) and adhesion factors (CD18, *PXN*).

As the prior analysis had been carried out on pooled clusters from all datasets, to be certain that results were not being preferentially skewed by one dataset, we separated the datasets again and compared the Pearson’s R of each dataset to random a gene selected by random number generation (*ARFIP2*), and present in all queried datasets.

All queried genes showed strong correlation across all four of the datasets: *C1orf21* (p = 0.0028), *STOM* (p = 0.0069), *S1PR5* (p = 0.0032), *PXN* (p = 0.0031), *PRF1* (p = 0.0072), and *CCL4* (p = 0.0032) (**Figure 3E**).

The same trend was visible when looking at the top correlated proteins: CD18 (*ITGB2*, p = 0.0018), CX3CR1 (p = 0.0064), CD16 (*FCGR3A*, p = 0.0025), CD244 (*SLAMF4*, p = 0.0047), CD45RA (p = 0.0090) and CD43 (*SPN*, p = 0.0018) (**Figure 3F**).

### Adhesion molecules are enriched among SH2D2A-interacting proteins in primary human PBMCs

We next investigated which proteins differentially expressed in SH2D2A^+^ cells were physically associated with SH2D2A. Primary human lymphocyte cultures were prepared with varying immune cell contents (summarised in **Figure 4A**), including unstimulated, PFA-fixed T cell blasts, and T cells stimulated with PV for 5 min. All cell cultures were expanded before being rested prior to activation and lysis.

**Figure 4.**
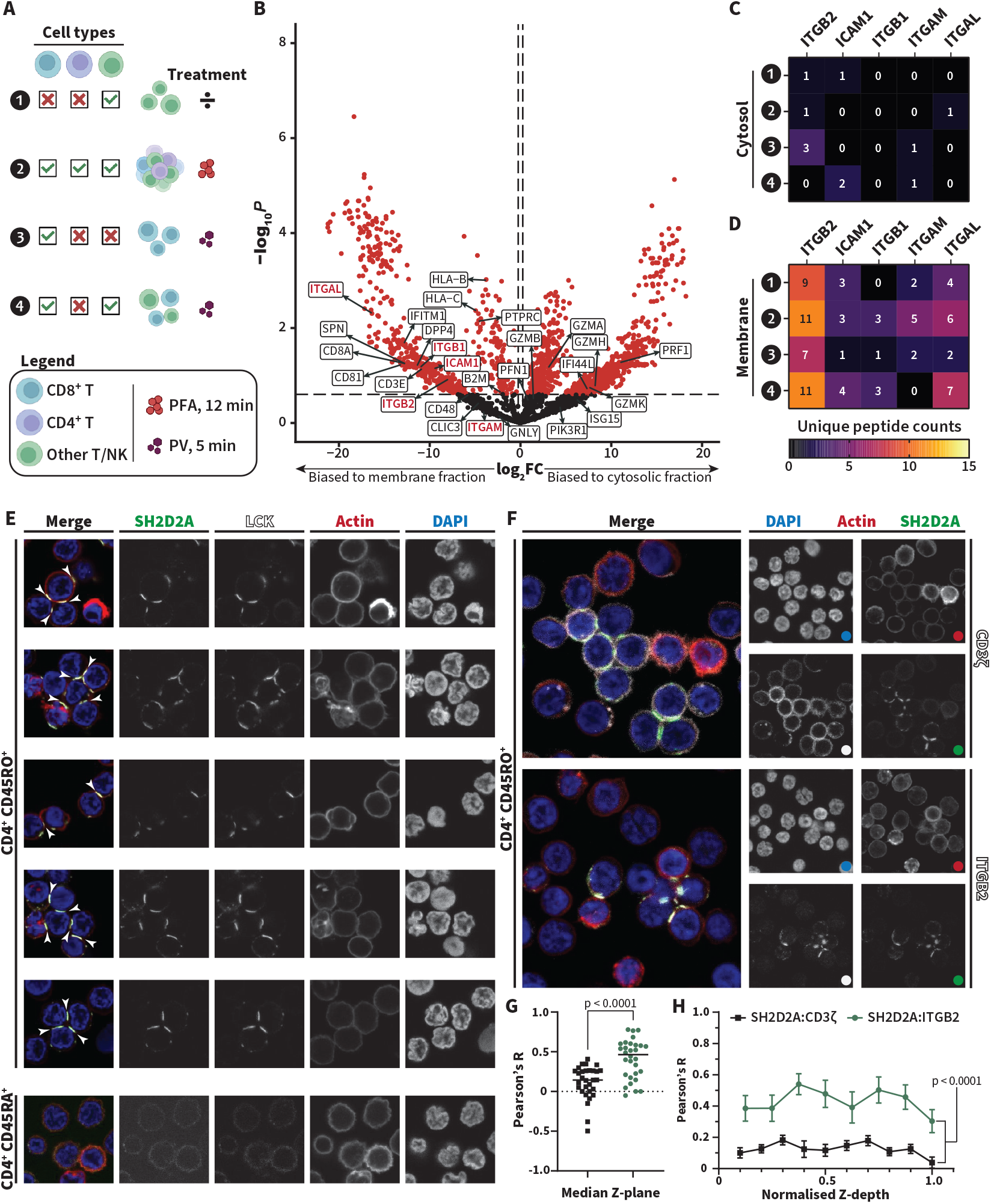
Adhesion molecules are enriched among SH2D2A-associated proteins, and SH2D2A colocalizes at T-T junctions with LCK and ITGB2, but not with CD3ζ. **(A)** Graphical summary of the samples used in mass spectrometry analysis, *(1)* PBMCs without CD4^+^ or CD8^+^ T cells, unstimulated; *(2)* full PBMCs with both CD4^+^ and CD8^+^ T cells, fixed with 4% PFA prior to lysis to preserve cell-cell contacts in the expanding blasts; *(3)* CD8^+^ T cells stimulated with PV for 5 min before lysis, and *(4)* PBMCs without CD4^+^ T cells, stimulated with PV for 5 min before lysis. Samples were lysed to isolate the cytosolic and membrane-bound fractions, and SH2D2A-associated proteins pulled down by anti-SH2D2A monoclonal antibody. **(B)** Volcano plot of proteins identified by mass spectrometry following pulldown of SH2D2A from the samples in **(A)**, proteins identified by differential gene expression or cluster correlation analysis are annotated, integrin family members and integrin-related proteins are marked (red). An unpaired, parametric, two-tailed Student’s *t*-test was used to determine statistical significance of each protein’s enrichment in the membrane (left) or cytosolic (right) fractions. **(C-D)** Unique peptide counts of the identified integrin family members and integrin-related proteins in the cytosol **(C)** and membrane **(D)** (…*) (*…) fractions. **(E)** Confocal microscopy of primary human CD4^+^ memory (CD45RO^+^, top) T cells stained for SH2D2A (green), LCK (white), actin (red) and the nucleus (blue). Areas of SH2D2A and LCK abundance at cell-cell junctions are marked (white arrows), a representative image of possible T-T interaction in naïve (CD45RA^+^, bottom) T cells is shown for comparison. Representative clusters from larger field images (n = 3, representative field images are shown in **Figure S3**). **(F)** confocal microscopy of CD4^+^ memory T cells stained for SH2D2A (green), actin (red), and the nucleus (blue), contrasting staining of the TCR-component CD3ζ (top, white) and the integrin ITGB2 (bottom, white). **(G)** Correlation analysis between SH2D2A and CD3ζ signals (SH2D2A:CD3ζ, black) and SH2D2A and ITGB2 signals (SH2D2A:ITGB2, green). Correlation analysis was performed on a per-cell basis, every dot represents one cell as defined by segmentation using the actin signal (n = 30 cells), statistical significance was determined by an unpaired, parametric, two-tailed Student’s *t*-test. **(H)** Comparison of correlation between SH2D2A:CD3ζ (black) and SH2D2A:ITGB2 (green) across Z-series, Z-depths of all individual cells were normalised to the number of Z-stacks in each cell, to account for varying cell sizes, each dot represents the average Pearson’s R of each cell at the given normalised Z-depth (n = 30 for SH2D2A:CD3ζ, n = 11 for SH2D2A:ITGB2) marked with the standard error of the mean (SEM). Statistical significance of the difference between the two lines was determined by ANCOVA using the model: *Pearson’s correlation coefficient ~ comparison* (*i*.*e*. SH2D2A:CD3ζ vs. SH2D2A:ITGB2) *+ z-depth*.

All samples were lysed to isolate first the cytosolic, then the membrane-associated proteome, and SH2D2A-associated proteins were pulled down using anti-SH2D2A antibody-coated beads. Mass spectrometry analysis revealed proteins preferentially associated with membrane (left) or cytosolic (right) fractions (**Figure 4B**).

The results, annotated with potential anchors from differential gene expression and cluster correlation analyses (**Figures 2–3**), showed enrichment of integrin-family and integrin-related proteins (**Figure 4B**, red). Tallying peptide counts in the cytosolic (Figure 4C) and membrane (**Figure 4D**) fractions, we found that SH2D2A pulldown of integrin family proteins was predominantly confined to the membrane fraction. ITGB2 (CD18) had the highest peptide count (9.5), followed by other integrin-related proteins, including ICAM1 (2.75), ITGB1 (1.75), ITGAM (2.24), and ITGAL (4.75).

### SH2D2A colocalizes at T-T junctions with LCK and ITGB2, but not with CD3ζ

Integrins mediate T-T interactions, along with other cell-cell interactions, all crucial for T cell activation, expansion, and differentiation^**41,42**^. Given SH2D2A’s association with ITGB2 (CD18), we examined its role in cell-cell interactions. In primary human CD4^+^ T cells, SH2D2A and its known interactor, LCK, accumulated at T-T junctions in memory CD45RO^+^ cells but not in naïve CD45RA^+^ cells (**Figure 4E**). Memory T cells also consistently exhibited more frequent T-T interactions (**Figure S4**).

As a known interactor of LCK, there was the possibility that SH2D2A was associated with an actively signalling TCR. To test whether this noted SH2D2A enrichment was linked to TCR signalling, we stained memory CD4^+^ CD45RO^+^ T cells for SH2D2A alongside either CD3ζ or ITGB2. SH2D2A co-localized with ITGB2 at T-T junctions but not with CD3ζ, which showed a pan-membrane distribution (**Figure 4F**). Pearson’s R correlation between SH2D2A and ITGB2 (0.41) was significantly higher than for SH2D2A and CD3ζ (0.12, p < 0.0001) (**Figure 4G**). Z-stack confocal analysis confirmed this pattern, with ANCOVA analysis showing significant correlation differences (p = 1.51 × 10^-9^) independent of Z-depth (p = 0.26) (**Figure 4H**).

## Discussion

At the outset of this study, we theorised that one could hypothetically work from the transcriptomic level upwards (thus “bottom-up”) to determine possible anchors—phenomena, cell states, and receptors—that could be used for further study of the protein in question. We have now carried out a trial analysis using a prototype bioinformatic approach based on this hypothesis and using the immune cell-enriched adaptor protein SH2D2A as a candidate gene. Through the application of this approach, we have been able to demonstrate its possible usefulness, managing to connect the presence of SH2D2A to cytotoxicity in various effector cells—*i*.*e*. CD8^+^ memory T cells, innate-like T cells (MAIT, γδT), and NK cells—as well as to cell-cell adhesion.

From here we have been able to expand our conclusions drawn from bioinformatic analysis to the laboratory, by connecting SH2D2A and its ligand LCK to T-T synapses, as well as to integrin-family proteins, especially to ITGB2—a molecule identified several times in our bioinformatic analysis. Indeed, SH2D2A has been previously linked to several adhesion or adhesion-adjacent phenomena in which integrins might play an important role. For example, SH2D2A has previously been implicated in the induction of polarity of CD4^+^ T cell synapses with antigen-presenting cells^**38**^, skewing their differentiation towards effector memory rather than central memory cells. Likewise, in Jurkat cells, lack of SH2D2A has shown to impact CXCL12-induced migration^**32**^, which was linked to SH2D2A’s interaction with ITK. An effect of SH2D2A on migration has been observed several times in Jurkat cells, and variously linked to effects downstream of laminin-binding protein (LBP)^**39**^ and the G protein subunit β^**40**^. Additionally, SH2D2A is a known regulator of cadherin junctions in lung epithelia, where it affects vascular permeability^**24**^.

Our work thus adds to this body of literature, and further suggests that the function of SH2D2A in immune cells may lie away from proximal TCR signalling, and closer to phenomena related to cell-cell adhesion. Whether the root phenomenon is T-T adhesion, migration, or the induction of synaptic polarity, remains to be seen. Importantly, while we have demonstrated a colocalisation of SH2D2A, LCK, and ITGB2 at T-T junctions, we have not demonstrated the physical interaction of these various molecules at these contact sites, and work continues to determine what SH2D2A is binding to within these junctions.

Further application of the prototype bottom-up approach as detailed in this paper to other immune proteins with uncertain function and few anchors could contribute greatly to the understanding of immune cell function. We have, however, for the time being only demonstrated the use of such an approach with a single gene, SH2D2A, and it remains to be seen whether such an analysis could prove useful for other protein cases.

## Supporting information

STAR Methods

## Abbreviations

ADT: antibody-dependent tag
ANCOVA: analysis of covariance
COVID-19: coronavirus disease 2019
DAPI: 4′,6-diamidino-2-phenylindole
FC: fold change
FDR: false discovery rate
HCD: higher-energy collisional dissociation
IF: immunofiuorescence
IJM: ImageJ macro
ITGB2: integrin β2
MAIT: mucosal-associated invariant T cell
MCL: Markov cluster
MS: mass spectrometry
NK: natural killer
PBMC: peripheral blood mononuclear cell
PCA: principal component analysis
PFA: paraformaldehyde
PLA: proximity ligation assay
PV: pervanadate
scRNA-seq: single cell RNA sequencing
SH2D2A: SH2 domain-containing 2A
TCR: T cell receptor
TSAd: T cell-specific adaptor protein
UMAP: uniform manifold approximation and projection
nUMI: normalised unique molecular identifier
VEGF: vascular endothelial growth factor

## Author contributions

Conceptualisation, B.C.G., P.B., and A.S.; Methodology, B.C.G; Software, B.C.G.; Formal Analysis, B.C.G.; Investigation, B.C.G; Resources, A.S.P., S.S., T.A.N. and A.L.; Data Curation, B.C.G.; Writing – original draft, B.C.G and A.S.; Writing – review & editing, B.C.G., H.C., P.B. and A.S; Visualisation, B.C.G; Supervision, A.S.P., S.P., P.B.; Funding Acquisition, A.S.P. and A.L.

## Acknowledgements

We would like to thank the MolMed Imaging Platform (MIP) at the Institute of Basic Medical Sciences, University of Oslo, for providing access to and training on relevant microscopes. Mass spectrometry-based proteomic analyses were performed by the Proteomics Core Facility, Department of Immunology, University of Oslo and Oslo University Hospital, which is supported by the Core Facilities program of the South-Eastern Norway Regional Health Authority. This core facility is also a member of the National Network of Advanced Proteomics Infrastructure, which is funded by the Research Council of Norway INFRASTRUKTUR program.

## References

1. Mosmann, T.R., and Coffman, R.L. (1989). TH1 and TH2 Cells: Different Patterns of Lymphokine Secretion Lead to Different Functional Properties. Annual Review of Immunology 7, 145–173. 10.1146/annurev.iy.07.040189.001045.

2. Boyman, O., and Sprent, J. (2012). The role of interleukin-2 during homeostasis and activation of the immune system. Nature Reviews Immunology 12, 180–190. 10.1038/nri3156.

3. Rochman, Y., Spolski, R., and Leonard, W.J. (2009). New insights into the regulation of T cells by γc family cytokines. Nature Reviews Immunology 9, 480–490. 10.1038/nri2580.

4. Cox, M.A., Kahan, S.M., and Zajac, A.J. (2013). Anti-viral CD8 T cells and the cytokines that they love. Virology 435, 157–169. 10.1016/j.virol.2012.09.012.

5. Akira, S., and Takeda, K. (2004). Toll-like receptor signalling. Nature Reviews Immunology 4, 499–511. 10.1038/nri1391.

6. Kumar, H., Kawai, T., and Akira, S. (2009). Pathogen recognition in the innate immune response. Biochemical Journal 420, 1–16. 10.1042/BJ20090272.

7. Geijtenbeek, T.B.H., and Gringhuis, S.I. (2009). Signalling through C-type lectin receptors: shaping immune responses. Nature Reviews Immunology 9, 465–479. 10.1038/nri2569.

8. Rehwinkel, J., and Gack, M.U. (2020). RIG-I-like receptors: their regulation and roles in RNA sensing. Nature Reviews Immunology 20, 537–551. 10.1038/s41577-020-0288-3.

9. Platnich, J.M., and Muruve, D.A. (2019). NOD-like receptors and inflammasomes: A review of their canonical and non-canonical signaling pathways. Archives of Biochemistry and Biophysics 670, 4–14. 10.1016/j.abb.2019.02.008.

10. Keating, S.E., Baran, M., and Bowie, A.G. (2011). Cytosolic DNA sensors regulating type I interferon induction. Trends in Immunology 32, 574–581. 10.1016/j.it.2011.08.004.

11. Gaud, G., Lesourne, R., and Love, P.E. (2018). Regulatory mechanisms in T cell receptor signalling. Nature Reviews Immunology 18, 485–497. 10.1038/s41577-018-0020-8.

12. Acuto, O., and Michel, F. (2003). CD28-mediated co-stimulation: a quantitative support for TCR signalling. Nature Reviews Immunology 3, 939–951. 10.1038/nri1248.

13. Patsoukis, N., Wang, Q., Strauss, L., and Boussiotis, V.A. Revisiting the PD-1 pathway. Science Advances 6, eabd2712. 10.1126/sciadv.abd2712.

14. Elgueta, R., Benson, M.J., De Vries, V.C., Wasiuk, A., Guo, Y., and Noelle, R.J. (2009). Molecular mechanism and function of CD40/CD40L engagement in the immune system. Immunological Reviews 229, 152–172. 10.1111/j.1600-065X.2009.00782.x.

15. Borowicz, P., Chan, H., Hauge, A., and Spurkland, A. (2020). Adaptor proteins: Flexible and dynamic modulators of immune cell signalling. Scandinavian Journal of Immunology 92, e12951. 10.1111/sji.12951.

16. Flynn, D.C. (2001). Adaptor proteins. Oncogene 20, 6270–6272. 10.1038/sj.onc.1204769.

17. Saba-El-Leil, M.K., Frémin, C., and Meloche, S. (2016). Redundancy in the World of MAP Kinases: All for One. Frontiers in Cell and Developmental Biology 4. 10.3389/fcell.2016.00067.

18. Sun, C., and Bernards, R. (2014). Feedback and redundancy in receptor tyrosine kinase signaling: relevance to cancer therapies. Trends in Biochemical Sciences 39, 465–474. 10.1016/j.tibs.2014.08.010.

19. June, C.H., Fletcher, M.C., Ledbetter, J.A., and Samelson, L.E. (1990). Increases in tyrosine phosphorylation are detectable before phospholipase C activation after T cell receptor stimulation. The Journal of Immunology 144, 1591–1599. 10.4049/jimmunol.144.5.1591.

20. Zhang, W., Sloan-Lancaster, J., Kitchen, J., Trible, R.P., and Samelson, L.E. (1998). LAT: The ZAP-70 Tyrosine Kinase Substrate that Links T Cell Receptor to Cellular Activation. Cell 92, 83–92. 10.1016/S0092-8674(00)80901-0.

21. Kolltveit, K.M., Granum, S., Aasheim, H.C., Forsbring, M., Sundvold-Gjerstad, V., Dai, K.Z., Molberg, O., Schjetne, K.W., Bogen, B., Shapiro, V.S., et al. (2008). Expression of SH2D2A in T-cells is regulated both at the transcriptional and translational level. Mol Immunol 45, 2380–2390. 10.1016/j.molimm.2007.11.005.

22. Dai, K.Z., Johansen, F.E., Kolltveit, K.M., Aasheim, H.C., Dembic, Z., Vartdal, F., and Spurkland, A. (2004). Transcriptional activation of the SH2D2A gene is dependent on a cyclic adenosine 5’-monophosphate-responsive element in the proximal SH2D2A promoter. J Immunol 172, 6144–6151. 10.4049/jimmunol.172.10.6144.

23. Sundvold, V., Torgersen, K.M., Post, N.H., Marti, F., King, P.D., Røttingen, J.A., Spurkland, A., and Lea, T. (2000). T cell-specific adapter protein inhibits T cell activation by modulating Lck activity. J Immunol 165, 2927–2931. 10.4049/jimmunol.165.6.2927.

24. Sun, Z., Li, X., Massena, S., Kutschera, S., Padhan, N., Gualandi, L., Sundvold-Gjerstad, V., Gustafsson, K., Choy, W.W., Zang, G., et al. (2012). VEGFR2 induces c-Src signaling and vascular permeability in vivo via the adaptor protein TSAd. J Exp Med 209, 1363–1377. 10.1084/jem.20111343.

25. Matsumoto, T., Bohman, S., Dixelius, J., Berge, T., Dimberg, A., Magnusson, P., Wang, L., Wikner, C., Qi, J.H., Wernstedt, C., et al. (2005). VEGF receptor-2 Y951 signaling and a role for the adapter molecule TSAd in tumor angiogenesis. EMBO J 24, 2342–2353. 10.1038/sj.emboj.7600709.

26. Wu, L.-W., Mayo, L.D., Dunbar, J.D., Kessler, K.M., Ozes, O.N., Warren, R.S., and Donner, D.B. (2000). VRAP Is an Adaptor Protein That Binds KDR, a Receptor for Vascular Endothelial Cell Growth Factor*. Journal of Biological Chemistry 275, 6059–6062. 10.1074/jbc.275.9.6059.

27. Granum, S., Sundvold-Gjerstad, V., Dai, K.Z., Kolltveit, K.M., Hildebrand, K., Huitfeldt, H.S., Lea, T., and Spurkland, A. (2006). Structure function analysis of SH2D2A isoforms expressed in T cells reveals a crucial role for the proline rich region encoded by SH2D2A exon 7. BMC Immunol 7, 15. 10.1186/1471-2172-7-15.

28. Granum, S., Andersen, T.C., Sørlie, M., Jørgensen, M., Koll, L., Berge, T., Lea, T., Fleckenstein, B., Spurkland, A., and Sundvold-Gjerstad, V. (2008). Modulation of Lck function through multisite docking to T cell-specific adapter protein. J Biol Chem 283, 21909–21919. 10.1074/jbc.M800871200.

29. Sundvold-Gjerstad, V., Granum, S., Mustelin, T., Andersen, T.C., Berge, T., Shapiro, M.J., Shapiro, V.S., Spurkland, A., and Lea, T. (2005). The C terminus of T cell-specific adapter protein (TSAd) is necessary for TSAd-mediated inhibition of Lck activity. Eur J Immunol 35, 1612–1620. 10.1002/eji.200425638.

30. Choi, Y.B., Kim, C.K., and Yun, Y. (1999). Lad, an adapter protein interacting with the SH2 domain of p56lck, is required for T cell activation. J Immunol 163, 5242–5249. 10.4049/jimmunol.163.10.5242.

31. Rajagopal, K., Sommers, C.L., Decker, D.C., Mitchell, E.O., Korthauer, U., Sperling, A.I., Kozak, C.A., Love, P.E., and Bluestone, J.A. (1999). RIBP, a novel Rlk/ Txk- and itk-binding adaptor protein that regulates T cell activation. J Exp Med 190, 1657–1668. 10.1084/jem.190.11.1657.

32. Berge, T., Sundvold-Gjerstad, V., Granum, S., Andersen, T.C.B., Holthe, G.B., Claesson-Welsh, L., Andreotti, A.H., Inngjerdingen, M., and Spurkland, A. (2010). T Cell Specific Adapter Protein (TSAd) Interacts with Tec Kinase ITK to Promote CXCL12 Induced Migration of Human and Murine T Cells. PLOS ONE 5, e9761. 10.1371/journal.pone.0009761.

33. Marti, F., Garcia, G.G., Lapinski, P.E., MacGregor, J.N., and King, P.D. (2006). Essential role of the T cell–specific adapter protein in the activation of LCK in peripheral T cells. Journal of Experimental Medicine 203, 281–287. 10.1084/jem.20051637.

34. Liu, C., Martins, A.J., Lau, W.W., Rachmaninoff, N., Chen, J., Imberti, L., Mostaghimi, D., Fink, D.L., Burbelo, P.D., Dobbs, K., et al. (2021). Time-resolved systems immunology reveals a late juncture linked to fatal COVID-19. Cell 184, 1836-1857.e1822. 10.1016/j.cell.2021.02.018.

35. Hao, Y., Hao, S., Andersen-Nissen, E., Mauck, W.M., Zheng, S., Butler, A., Lee, M.J., Wilk, A.J., Darby, C., Zager, M., et al. (2021). Integrated analysis of multimodal single-cell data. Cell 184, 3573-3587.e3529. 10.1016/j.cell.2021.04.048.

36. Szklarczyk, D., Kirsch, R., Koutrouli, M., Nastou, K., Mehryary, F., Hachilif, R., Gable, A.L., Fang, T., Doncheva, Nadezhda T., Pyysalo, S., et al. (2023). The STRING database in 2023: protein–protein association networks and functional enrichment analyses for any sequenced genome of interest. Nucleic Acids Research 51, D638–D646. 10.1093/nar/gkac1000.

37. Van Dongen, S. (2008). Graph Clustering Via a Discrete Uncoupling Process. SIAM Journal on Matrix Analysis and Applications 30, 121–141. 10.1137/040608635.

38. Abrahamsen, G., Sundvold-Gjerstad, V., Habtamu, M., Bogen, B., and Spurkland, A. (2018). Polarity of CD4+ T cells towards the antigen presenting cell is regulated by the Lck adapter TSAd. Scientific Reports 8, 13319. 10.1038/s41598-018-31510-6.

39. Park, E., Choi, Y., Ahn, E., Park, I., and Yun, Y. (2009). The adaptor protein LAD/TSAd mediates laminin-dependent T cell migration via association with the 67 kDa laminin binding protein. Experimental & Molecular Medicine 41, 728–736. 10.3858/emm.2009.41.10.079.

40. Park, D., Park, I., Lee, D., Choi, Y.B., Lee, H., and Yun, Y. (2007). The adaptor protein Lad associates with the G protein β subunit and mediates chemokine-dependent T-cell migration. Blood 109, 5122–5128. 10.1182/blood-2005-10-061838.

41. Gérard, A., Khan, O., Beemiller, P., et al. (2013). Secondary T cell-T cell synaptic interactions drive the differentiation of protective CD8+ T cells. Nat Immunol 14, 356–363. 10.1038/ni.2547.

42. Sabatos, C.A., Doh, J., Chakravarti, S., Friedman, R.S., Pandurangi, P.G., Tooley, A.J., and Krummel, M.F. (2008). A Synaptic Basis for Paracrine Interleukin-2 Signaling during Homotypic T Cell Interaction. Immunity 29, 238–248. 10.1016/j.immuni.2008.05.017.

