## Supplementary material for "A bioinformatic workflow to facilitate the study of less-understood proteins: the case of SH2D2A": STAR Methods

### Key Resource Table

| REAGENT or RESOURCE | SOURCE | IDENTIFIER |
|---|---|---|
| **Antibodies** | | |
| Anti-CD18 (polyclonal) | Nordic Biosite | LS-C117844 |
| Anti-CD247 (BL-336-1B2) | ThermoFisher | A700-017 |
| Anti-LCK (Y123) | Abcam | ab32149 |
| Anti-SH2D2A (OTI3C7) | Origene | TA504372S |
| Goat anti-Mouse IgG (H+L), Alexa Fluor™ 488 | ThermoFisher | A-11029 |
| Goat anti-Rabbit IgG (H+L), Alexa Fluor™ 647 | ThermoFisher | A-21245 |
| **Biological samples** | | |
| Buffy coat | Norwegian Blood Bank | S-09325b |
| **Chemicals, peptides, and recombinant proteins** | | |
| 1,4-Dithiothreitol (DTT) | Merck | 111474 |
| Acetonitrile | Merck | 100030 |
| Ammonium hydrogen carbonate ($NH_4HCO_3$) | Merck | 101131 |
| Bovine serum albumin (BSA) | Bio-Rad | 805095 |
| DAPI | ThermoFisher | 62248 |
| Digitonin | Merck | 300410 |
| Fetal bovine serum (FBS) | ThermoFisher | A5256701 |
| Formic acid | Merck | 100264 |
| GlutaMAX™ Supplement | ThermoFisher | 35050061 |
| HEPES 1M | ThermoFisher | 15630-056 |
| Hexylene glycol | Merck | 820819 |
| Human IL-2 recombinant protein | ThermoFisher | 200-02 |
| Hydrogen peroxide ($H_2O_2$) | Merck | 216763 |
| Iodoacetamide | Merck | I1149 |
| Minimal essential media non-essential amino acids 100X | ThermoFisher | 11140-035 |
| Nonidet P-40 | Merck | 56741 |
| Octyl β-D-glucopyranoside | Merck | O8001 |
| Paraformaldehyde | Merck | P6148 |
| Penicillin-streptomycin | ThermoFisher | 15140-122 |
| Phalloidin Alexa Fluor™ 546 | ThermoFisher | A22283 |
| ProteaseMAX™ Surfactant | Promega | V207A |
| RPMI 1640 | ThermoFisher | 21875-091 |
| SIGMAFAST™ Protease Inhibitor cocktail | Merck | S8830 |
| SlowFade™ Gold Antifade Mountant | ThermoFisher | S36936 |
| Sodium chloride (NaCl) | Merck | 1.06404 |
| Sodium fluoride (NaF) | Merck | 71518 |
| Sodium orthovanadate ($Na_3VO_4$) | Merck | S6508 |
| Sodium pyruvate 100 nM | ThermoFisher | 11360-039 |
| Tris Base | Merck | 648310-M |
| Triton X-100 | Merck | T8787 |
| Tween® 20 | VWR | 28829.296 |
| β-mercaptoethanol | Merck | M3148 |
| **Critical commercial assays** | | |
| Dynabeads® FlowComp™ Human CD45RA | ThermoFisher | 11368D |

| REAGENT or RESOURCE | SOURCE | IDENTIFIER |
|---|---|---|
| Dynabeads™ CD4 Positive Isolation Kit | ThermoFisher | 11331D |
| Dynabeads™ CD8 Positive Isolation Kit | ThermoFisher | 11333D |
| Dynabeads™ Human T-Activator CD3/CD28 | ThermoFisher | 11131D |
| Dynabeads™ Protein G for Immunoprecipitation | ThermoFisher | 10003D |
| In Situ PLA® Detection Reagents FarRed | Merck | DUO92008 |
| In Situ PLA® Probe Anti-Mouse MINUS | Merck | DUO92004 |
| In Situ PLA® Probe Anti-Rabbit PLUS | Merck | DUO92002 |
| **Software and algorithms** | | |
| Fiji | Open Source | 2.9.0 |
| Prism | GraphPad | 10.4.1 |
| MaxQuant | Max-Planck Institute of Biochemistry | 1.6.17.0 |
| R | The R Project | 4.4.2 |
| RStudio | Posit PBC | 2024.12.0 |
| STRING | String Consortium | 12.0 |
| Zen | Carl Zeiss Imaging | 3.11 |
| **Other** | | |
| 8-well glass-bottomed Lab-Tek® Chamber Slides™ | ThermoFisher | 177402 |
| C18 Empore™ Extraction Disk | Varian | 13-110-018 |

### Experimental model and study participant details

#### Peripheral blood mononuclear cells (PBMCs) from human buffy coat donations

Buffy coats sourced from anonymized donations to the Norwegian Blood Bank (under project **S-09325b**) were used as a source of peripheral blood mononuclear cells from which naïve ($CD45RA^+$) and memory ($CD45RO^+$) $CD8^+$ and $CD4^+$ T cells were isolated. Due to anonymization, information on sex, age, HLA-type, etc., was not available.

#### Sourced public datasets

This study makes use of four publicly available datasets containing data from human patients and/or healthy controls. Information on the study participants and experimental models can be found in the original publications of each dataset, as listed below:

1. Liu C, Martins AJ, Lau WW, *et al.* Time-resolved systems immunology reveals a late juncture linked to fatal COVID-19. *Cell.* 2021;**184**(7):1836-1857.e22. 10.1016/j.cell.2021.02.018
2. Hao Y, Hao S, Andersen-Nissen E, *et al.* Integrated analysis of multimodal single-cell data. *Cell.* 2021;**184**(13):3573-3587.e29. 10.1016/j.cell.2021.04.048
3. Wilk AJ, Rustagi A, Zhao NQ, *et al.* A single-cell atlas of the peripheral immune response in patients with severe COVID-19. *Nat Med.* 2020;**26**(7):1070-1076. 10.1038/s41591-020-0944-y
4. Zhang JY, Wang XM, Xing X, *et al.* Single-cell landscape of immunological responses in patients with COVID-19. *Nat Immunol.* 2020;**21**(9):1107-1118. 10.1038/s41590-020-0762-x

### Method details

#### Data acquisition

All scRNA-seq datasets were downloaded from publicly available repositories. The dataset from Liu *et al.* (2021)[1] was downloaded as paired expression matrices and metadata files for the adaptive and innate immune cells from UCSC Cell Browser. Hao *et al.* (2021)[2] was originally downloaded from the New York Genome Center and used to extrapolate values from a 228-antibody panel antibody-dependent tag (ADT) assay of immune-relevant surface proteins onto the Liu *et al.* dataset using *Seurat*[3] – this can otherwise be done using the R package *Azimuth* and its *pbmcref* dataset. The datasets from Wilk *et al.* (2020)[4] and Zhang *et al.* (2020)[5] were downloaded from UCSC Cell Browser. Only healthy donors from the Wilk *et al.* dataset were used in this study. The dataset from Zhang *et al.* was further split by patient condition, and the data subsets consisting of healthy donors and convalescent patients used for correlation analysis.

#### Preparation of the mapped Liu *et al.* dataset

Data from Liu *et al.* (2021)[1] was downloaded as two separate datasets: the first consisting of adaptive immune cells, the second consisting of innate immune cells. Cells annotated as $CD4^+$ T, $CD8^+$ T and Other T cells were isolated from the adaptive immune cell dataset, and cells annotated as NK cells were isolated from the innate immune cell dataset. The T and NK cell datasets were then combined, and the resulting merged dataset analysed as per standard scRNA-seq analysis with *Seurat*. Briefly: data was normalised, a principal component analysis (PCA) run to reduce dimensionality, and the result plotted via uniform manifold approximation and projection (UMAP) to produce the modified Liu *et al.* (MOD-Liu) dataset consisting of T and NK cells and data from the original scRNA-seq assay. The peripheral blood mononuclear cell (PBMC) reference dataset from Hao *et al.* (2021)[2], consisting of paired data from an scRNA-seq assay and an antibody-dependent tag (ADT, CiteSeq)[6] assay of 228 immune-relevant TotalSeq™ antibodies, was then used to extrapolate ADT data onto the MOD-Liu dataset, producing the mapped Liu *et al.* dataset (MAP-Liu) consisting of paired scRNA-seq assay data and extrapolated ADT assay data. An overview of the different forms of the Liu *et al.* dataset used in this study can be found in **Table S1**.

### Isolation of primary $CD45RA^+$ and $CD45RO^+$ $CD4^+$ and $CD8^+$ T cells from buffy coat

Human buffy coats were sourced from the Norwegian Blood Bank under project number S-09325b. $CD4^+$ and $CD8^+$ T cells were isolated from buffy coats by Dynabeads® CD4 Positive or CD8 Positive Isolation Kits (both ThermoFisher) as per the manufacturer's protocol. Isolated cells were then used for downstream assays, cultured, or further separated into naïve and memory portions. Dynabeads® FlowComp™ Human CD45RA kit (Invitrogen) was used to separate cultures into naïve ($CD45RA^+$) and memory ($CD45RO^+$) portions as per the manufacturer's protocol. The bead-free cells were immediately used for proximity ligation assay or immunofluorescence.

### Culture and expansion of primary human T cells

Isolated primary human $CD4^+$ and $CD8^+$ T cells were cultured in RPMI 1640 supplemented with 10% FBS, 1% penicillin-streptomycin, 1X Minimum Essential Medium (MEM) Non-Essential Amino Acids (NEAA), 10 mM HEPES buffer, 1 mM sodium pyruvate, 1X GlutaMAX™ (all ThermoFisher) and 40 U/mL human recombinant IL-2 (ThermoFisher). Expansion of human T cell cultures was accomplished by adding 25 µL Dynabeads™ Human T-Activator CD3/CD28 beads (ThermoFisher) to the culture medium per $1 \times 10^6$ T cells, followed by on-bead incubation in culture for up to 7 days. Cells were detached from the beads by pipetting the culture solution up and down 10 times, and beads were then removed by a magnet. Cells were then allowed to rest overnight before cultures were harvested and immediately used for downstream applications.

### Stimulation of primary T cell cultures

To simulate TCR stimulation prior to proximity ligation assay, primary human $CD45RA^+$ and $CD45RO^+$ $CD4^+$ and $CD8^+$ T cells were activated with magnetic Dynabeads™ Human T-Activator CD3/CD28 beads (ThermoFisher) by incubation in phosphate buffered saline (PBS) at 37˚C for 5 min in a heat bath before activation was stopped by addition of an excess of ice-cold PBS. Cells were then detached from the beads by pipetting the culture solution up and down 10 times, and beads were then removed by magnet. For mass spectrometry analysis, sodium orthovanadate ($Na_3VO_4$, Merck) was used to generate pervanadate (PV) and treat cells to induce a state of hyperphosporylation through inhibition of phosphatase activity[7]. To this end, 100 mM $Na_3VO_4$ was diluted 1:10 with 1% $H_2O_2$ (Merck), then brought to a final concentration of 100 µM in the samples to be treated followed by incubation for 5 min at 37˚C. Treatment was stopped by addition of an excess of ice-cold PBS. In all cases, incubation in PBS for 5 min at 37˚C in a heat bath was used as a control.

### Antibodies and counterstains

The following antibodies were used: anti-SH2D2A (clone OTI3C7, Origene), anti-LCK (clone Y123, Abcam), anti-CD247 (clone BL-336-1B2, ThermoFisher), anti-CD18 (polyclonal, Nordic Biosite). The following secondary antibodies were used: Alexa Fluor 488 (AF488)-conjugated anti-Mouse IgG1 and AF647-conjugated anti-Rabbit IgG (H+L) (both ThermoFisher). The following molecules were used for counterstaining: DAPI (DNA stain) and AF546-conjugated phalloidin (actin stain, both ThermoFisher).

### Immunofluorescence

Cells were allowed to adhere to 8-well, glass-bottomed Lab-Tek® Chamber Slides™ (ThermoFisher) in PBS for 45 min at 37˚C and 5% $CO_2$, as described elsewhere[8], followed by fixation with 4% paraformaldehyde (PFA, Merck) in PBS at room temperature (RT) for 12 min. Cells were washed twice gently with PBS and permeabilized with 0.1% Triton X-100 (Merck) in PBS for 15 min at RT, followed by two gentle washes with PBS. Cells were incubated with 1% bovine serum albumin (BSA, Bio-Rad) in

PBS for 1 hr at RT. Plastic chambers were then removed, and cells were incubated with primary unconjugated antibody diluted in 1% BSA in PBS for 1 hr at RT. Following primary staining, cells were washed twice gently with 1% BSA in PBS. For secondary staining for IF, cells were incubated for 1 hr in the dark at RT with fluorophore-conjugated secondary antibodies. Cells were counterstained with DAPI and AF546-conjugated phalloidin diluted 1:3000 and 1:40, respectively, in 1% BSA in PBS for 15 min in the dark at RT, washed twice with 1% BSA in PBS, before being mounted with SlowFade™ Gold (ThermoFisher) antifade reagent. Mounted slides were then sealed with nail polish, and fluorescent signals allowed to mature at –20°C overnight before imaging by confocal microscopy.

### Proximity ligation assay (PLA)

Proximity ligation assay was performed similarly to IF, based on protocols described elsewhere[9,10], with cells instead permeabilized with 0.5% Triton X-100 in PBS for 15 min at RT. Primary antibody incubation for PLA occurred overnight at 4°C in a humidity chamber. PLA was accomplished using the following Merck Duolink® reagents: In Situ PLA® Probe Anti-Mouse MINUS, Anti-Rabbit PLUS, and In Situ Detection Reagents FarRed. The PLA reaction was carried out as per the manufacturer's instructions. Slides were then counterstained and mounted as detailed above for the procedure for immunofluorescence.

### Confocal microscopy and image processing

Microscopy slides were imaged on a Zeiss LSM 710 confocal microscope at 63x magnification. Z-series were taken of all images, with the lowest Z-slice beginning just above the cells' contact point with the glass slide, and the highest Z-slice just before the end of the tallest cell. Captured images were pre-processed in Zen (Carl Zeiss Imaging) before being loaded into Fiji[11]. Representative images were then selected for each image and Z-slice, and Fiji was used to split and export each colour channel as a separate image.

### Lysis and SH2D2A pulldown from primary human PBMCs

Expanded T cells were harvested from culture, spun down at 400g at RT for 8 min, and resuspended in 10 mL PBS. Cells were counted using a TC20 Automated Cell Counter (Bio-Rad), and cell numbers equalized. The following cell cultures were used: (1) PBMCs without $CD4^+$ or $CD8^+$ T cells, unstimulated; (2) full PBMCs with both $CD4^+$ and $CD8^+$ T cells, fixed with 4% PFA prior to lysis to preserve cell-cell contacts in the expanding blasts; (3) $CD8^+$ T cells stimulated with PV for 5 min before lysis, and (4) PBMCs without $CD4^+$ T cells stimulated with PV for 5 min before lysis. In the cases of PV treatment, stimulation was stopped by the addition of an excess of ice-cold PBS. A graphic summary of the various activating conditions included in this experiment is available in **Figure 4A**. All samples were then spun down at 400g at RT for 5 min and supernatants removed by suction. All samples were then lysed by resuspension in 700 µL per $10^6$ cells of digitonin lysis buffer (DG-LB) of 150 mM NaCl, 50 nM HEPES (pH 7.4), 1 M hexylene glycol, 1X SIGMA*FAST*™ Protease Inhibitor cocktail (Merck), 1 mM $Na_3VO_4$, 25 mM NaF, and 150 µg/mL digitonin in distilled water. The lysates were incubated on a rotating wheel for 10 min at 4°C to release the cytosolic fractions of the cells. To collect the cytosolic proteome, samples were then spun down for 10 min at 2,000g and 4°C and the supernatant removed. The remaining insoluble fraction, representing both the membrane and nucleus, was resuspended in 700 µL per $10^6$ cells of Nonidet P-40 lysis buffer (NP-LB) of 40 nM Tris (pH 7.5), 1X SIGMA*FAST*™ Protease Inhibitor cocktail, 1 mM $Na_3VO_4$, 25 mM Nonidet P-40, 1X octyl β-D-glucopyranoside (Merck) in distilled water. To release membrane bound proteins, samples were incubated on ice for 30 min, and vortexed every 10 min. Following lysis, all samples were spun for 10 min at 13,000g at 4°C to eliminate debris and remove the nuclei, in the case of the membrane samples. In order to pull-down proteins associated with SH2D2A, Dynabeads™ Protein G beads (ThermoFisher) were pre-washed three times with 0.01% Tween® 20 and then coated with anti-SH2D2A monoclonal antibody (clone OTI3C7) by incubation on a rotating wheel for ≥1 hr. Beads were then washed three times in their respective lysis buffer (DG-LB or NP-LB, with 0.2X SIGMA*FAST*™ Protease Inhibitor cocktail instead of 1X). All lysates were incubated with the antibody-loaded protein G beads on a rotating wheel for 1 hr at 4°C. SH2D2A-interacting proteins were isolated by magnetic isolation of the protein G beads. Non-specific protein interactions were removed first by washing the magnetic beads three times with wash buffer, transferring the beads to new tubes, and then washing 3 times with PBS. Isolated beads with captured proteins were then digested prior to mass spectrometry.

### Protein digestion and mass spectrometry

Beads with pulled down protein were resuspended in 10 µl 0.2% ProteaseMAX™ Surfactant (Promega) in 50 mM $NH_4HCO_3$ (Merck) and were further diluted in 100 µl 50 mM $NH_4HCO_3$. Cysteines were reduced in 5 mM DTT (Merck) at 56°C for 30 min and alkylated in 15 mM iodoacetamide (Merck) for 20 min at RT before overnight digestion by trypsin (1 µg, Promega) at 37°C. The resulting peptides were desalted by home-made STAGE tip C18 columns prepared by stacking three layers of C18 Empore™ Extraction Disk (Varian) into 200 µL pipette tips. Desalted peptide samples were analysed by EASY-nLC™ 1000 liquid chromatography system connected to a Q Exactive™ Plus Hybrid Quadrupole-Orbitrap™ Mass Spectrometer equipped with an EASY-Spray™ ion source (all ThermoFisher). An EASY-Spray™ column (C18, 2 µm beads, 100 Å, 75 µm inner diameter, ThermoFisher) capillary of 25 cm bed length was used for liquid chromatography separation. The flow rate used was 0.3 µL/min, and the solvent gradient was 2-7% for 5 min then 30% for 60 min. The column was finally washed in 90% B wash for 20 min. Solvent A was aqueous 0.1% formic acid, whereas solvent B was 100% acetonitrile in 0.1% formic acid. Column temperature was kept at 60˚C. The mass spectrometer was operated in data-dependent mode to automatically switch between MS and MS/MS acquisition. Full scans of MS spectra (m/z 300-1 750) were acquired in the Orbitrap with resolution R = 60,000 at m/z 200 (after accumulation to a target of 3,000,000 ions in the quadrupole), allowing sequential isolation of the most intense multiply-charged ions, up to ten, depending on signal intensity, for fragmentation on the HCD cell using high-energy collision dissociation at a target value of 100,000 charges or maximum acquisition time of 20 ms. MS/MS scans were collected at a resolution of 15,000 at the Orbitrap cell. Target ions already selected for MS/MS were dynamically excluded for 30 s. General mass spectrometry conditions were: electrospray voltage set to 2.1 kV, no sheath and auxiliary gas flow, heated capillary temperature of 250˚C, normalised HCD collision energy 25%.

### Quantification and statistical analysis

#### Analysis of the mapped Liu *et al.* dataset

Bottom-up analysis of the MAP-Liu dataset was performed with the help of *Seurat* in R and followed two main routes: *(1)* differential expression analysis within manually annotated cell types, and *(2)* correlation analysis between high-resolution clusters. Briefly, classic differential expression analysis was performed for each cell type contrasting the *SH2D2A*$^+$ and *SH2D2A*$^-$ fractions of each cell type. Identified genes and proteins were then checked for re-occurrence across several cell types, and STRING[12] used to visualise the network of the top 25 most upregulated genes in the cell type with the highest number of differentially expressed elements, naïve CD8$^+$ T cells. The correlation analysis between cell clusters used higher resolution during the *Seurat* clustering step to increase the number clusters and then averaged the expression of every gene and protein in each cluster, producing a pseudo-bulk expression value for each cluster for each gene and protein. The expression of all genes and proteins were then correlated to that of SH2D2A to determine elements which had a similar expression pattern, reporting only elements with a Pearson's $R^2 > 0.6$. To control for dataset specific artefacts in MAP-Liu, we sourced also data from Wilk *et al.* (2020)[4] and Zhang *et al.* (2020)[5]. To check for the specificity of the correlation, we selected a gene by random number generation (*ARFIP2*) to serve as a control for random correlation.

#### PLA image analysis

A custom ImageJ macro (IJM) script was used for PLA analysis, briefly: masks were made of the PLA signal, nucleus, and membrane channels (of the median Z-slice). The PLA signal mask was then simplified into discrete ellipses to combine converged signals. The raw count of PLA signal ellipses was then divided by the maximum number of nuclei across all Z-slices in an image. Concomitantly, the membrane mask of the median Z-slice was used to identify discrete cells and used to summarise the pixel intensity of the PLA signal within each cell. The membrane mask was also used to iterate through each cell in each picture to determine the per cell values of PLA intensity and PLA blob counts. A depiction of the general workflow of this script is available in **Figure S4**.

#### Analysis of mass spectrometry output

Raw mass spectrometry output was submitted to MaxQuant (v1.6.17.0) for protein identification and quantification. Parameters were set as follows: carbamidomethylation as fixed modification and protein N-acetylation, methionine oxidation and phosphorylation (STY) as variable modifications. First search error window of 20 ppm and main search error of 4.5 ppm. Trypsin without proline restriction enzyme option was used, with two allowed miscleavages. Minimal unique peptides were set to 1, and FDR allowed was 0.01 (1%) for peptide and protein identification. UniProt was used to map protein IDs to peptide sequences (accessed 2022). Statistical analysis of MaxQuant output was done in R, contrasting protein abundancies in all cytosolic fractions versus all membrane fractions. Unique peptide counts for identified integrin family members and integrin-associated proteins were exported directly from the MaxQuant output and tallied in Prism.

### Statistical analysis

R and GraphPad were used for statistical analysis. Wilcoxon signed-rank test with Bonferroni correction was used for differential gene expression analysis, reporting results with both positive and negative fold changes (FCs), and featuring an FC cutoff of ± 0.25. Linear correlation analysis was used to produce Pearson's correlation coefficients (Pearson's R). A paired, two-tailed parametric Student's *t*-test was used to determine consensus between correlation coefficients from different datasets between *SH2D2A* and *ARFIP2* (a randomly selected gene). Unpaired, two-tailed parametric Student's *t*-tests were used to determine significant hits from mass spectrometry, and test if differences between the SH2D2A:CD3ζ and SH2D2A:ITGB2 correlations were statistically different. Analysis of covariance (ANCOVA) was used to determine if differences in Pearson's R between SH2D2A:CD3ζ and SH2D2A:ITGB2 resulted from different Z-slices in the confocal images, or from the different comparisons themselves, using the model: *Pearson's correlation coefficient ~ comparison* (i.e. SH2D2A:CD3ζ vs. SH2D2A:ITGB2) + *Z-depth*. An α of 0.05 was use for all statistics, and p values reported as raw numbers.

### Code and data availability

All analytical work was carried out using pre-existing algorithms and packages, and no novel algorithms or packages were developed for the purpose of this study. The mass spectrometry proteomics data have been deposited to the ProteomeXchange Consortium via the PRIDE partner repository with the dataset identifier PXD061437. The scripts for the bioinformatic workflow detailed in this paper and the IJM analysis script for PLA ouput images are available on Mendley Data under the DOI: 10.17632/6fry5p2zkh.1. All scRNA-seq datasets were downloaded from publicly available repositories and are thus free to access. A summary of the datasets used in this study is available in **Table S2**.

### Additional resources

UCSC Cell Browser: https://cells.ucsc.edu/

The New York Genome Center: https://atlas.fredhutch.org/nygc/

### References


1. Liu, C., Martins, A.J., Lau, W.W., Rachmaninoff, N., Chen, J., Imberti, L., Mostaghimi, D., Fink, D.L., Burbelo, P.D., Dobbs, K., *et al.* (2021). Time-resolved systems immunology reveals a late juncture linked to fatal COVID-19. *Cell* **184**, 1836-1857.e1822. 10.1016/j.cell.2021.02.018.
2. Hao, Y., Hao, S., Andersen-Nissen, E., Mauck, W.M., Zheng, S., Butler, A., Lee, M.J., Wilk, A.J., Darby, C., Zager, M., *et al.* (2021). Integrated analysis of multimodal single-cell data. *Cell* **184**, 3573-3587.e3529. 10.1016/j.cell.2021.04.048.
3. Hao, Y., Stuart, T., Kowalski, M.H., Choudhary, S., Hoffman, P., Hartman, A., Srivastava, A., Molla, G., Madad, S., Fernandez-Granda, C., and Satija, R. (2023). Dictionary learning for integrative, multimodal and scalable single-cell analysis. *Nature Biotechnology*. 10.1038/s41587-023-01767-y.
4. Wilk, A.J., Rustagi, A., Zhao, N.Q., Roque, J., Martínez-Colón, G.J., McKechnie, J.L., Ivison, G.T., Ranganath, T., Vergara, R., Hollis, T., *et al.* (2020). A single-cell atlas of the peripheral immune response in patients with severe COVID-19. *Nature Medicine* **26**, 1070-1076. 10.1038/s41591-020-0944-y.
5. Zhang, J.-Y., Wang, X.-M., Xing, X., Xu, Z., Zhang, C., Song, J.-W., Fan, X., Xia, P., Fu, J.-L., Wang, S.-Y., *et al.* (2020). Single-cell landscape of immunological responses in patients with COVID-19. *Nature Immunology* **21**, 1107-1118. 10.1038/s41590-020-0762-x.
6. Stoeckius, M., Hafemeister, C., Stephenson, W., Houck-Loomis, B., Chattopadhyay, P.K., Swerdlow, H., Satija, R., and Smibert, P. (2017). Simultaneous epitope and transcriptome measurement in single cells. *Nature Methods* **14**, 865-868. 10.1038/nmeth.4380.
7. Zick, Y., and Sagi-Eisenberg, R. (1990). A combination of H2O2 and vanadate concomitantly stimulates protein tyrosine phosphorylation and polyphosphoinositide breakdown in different cell lines. *Biochemistry* **29**, 10240-10245. 10.1021/bi00496a013.
8. Tsang, M., Gantchev, J., Ghazawi, F.M., and Litvinov, I.V. (2017). Protocol for adhesion and immunostaining of lymphocytes and other non-adherent cells in culture. *BioTechniques* **63**, 230-233. 10.2144/000114610.
9. Alam, M.S. (2018). Proximity Ligation Assay (PLA). *Current Protocols in Immunology* **123**, e58. 10.1002/cpim.58.
10. Hegazy, M., Cohen-Barak, E., Koetsier, J.L., Najor, N.A., Arvanitis, C., Sprecher, E., Green, K.J., and Godsel, L.M. (2020). Proximity Ligation Assay for Detecting Protein-Protein Interactions and Protein Modifications in Cells and Tissues in Situ. *Current Protocols in Cell Biology* **89**, e115. 10.1002/cpcb.115.
11. Schindelin, J., Arganda-Carreras, I., Frise, E., Kaynig, V., Longair, M., Pietzsch, T., Preibisch, S., Rueden, C., Saalfeld, S., Schmid, B., *et al.* (2012). Fiji: an open-source platform for biological-image analysis. *Nature Methods* **9**, 676-682. 10.1038/nmeth.2019.
12. Szklarczyk, D., Kirsch, R., Koutrouli, M., Nastou, K., Mehryary, F., Hachilif, R., Gable, A.L., Fang, T., Doncheva, Nadezhda T., Pyysalo, S., *et al.* (2023). The STRING database in 2023: protein–protein association networks and functional enrichment analyses for any sequenced genome of interest. *Nucleic Acids Research* **51**, D638-D646. 10.1093/nar/gkac1000.
