## Supplemental Material for "A bioinformatic workflow to facilitate the study of less-understood proteins: the case of SH2D2A"

### Supplementary information

#### TABLE S1. Variations of the Liu et al. dataset used in this study.

| Alias | Name | Cell types | Assays | Description |
|---|---|---|---|---|
| Liu *et al.* (adaptive) | Liu *et al.* (adaptive) | Adaptive immune cells | RNA | All adaptive immune cells from the original publication. |
| Liu *et al.* (innate) | Liu *et al.* (innate) | Innate immune cells | RNA | All innate immune cells from the original publication. |
| MOD-Liu | Modified Liu *et al.* | T and NK cells | RNA | Dataset combining the T and NK cells from the above 2 datasets. |
| MAP-Liu | Mapped Liu *et al.* | T and NK cells | RNA + ADT | Extrapolation of ADT assay values for immune surface proteins from Hao *et al.* onto the MOD-Liu dataset. |

#### TABLE S2. Datasets used in this study.

| Name | Source | PMID | Type | Content |
|---|---|---|---|---|
| Liu *et al.* | UCSC Cell Browser | 33713619 | RNA | scRNA-seq of PBMCs isolated from COVID-19 patients and healthy donors. |
| Hao *et al.* (*pbmcref*) | New York Genome Center, *Azimuth* (R package) | 34062119 | RNA + ADT | Reference PBMC dataset containing a 228-antibody ADT assay |
| Wilk *et al.* | UCSC Cell Browser | 32514174 | RNA | scRNA-seq of PBMCs isolated from COVID-19 patients and healthy donors. |
| Zhang *et al.* | UCSC Cell Browser | 32788748 | RNA | scRNA-seq of PBMCs isolated from patients with moderate and severe COVID-19, from convalescent COVID-19 patients, and healthy donors. |

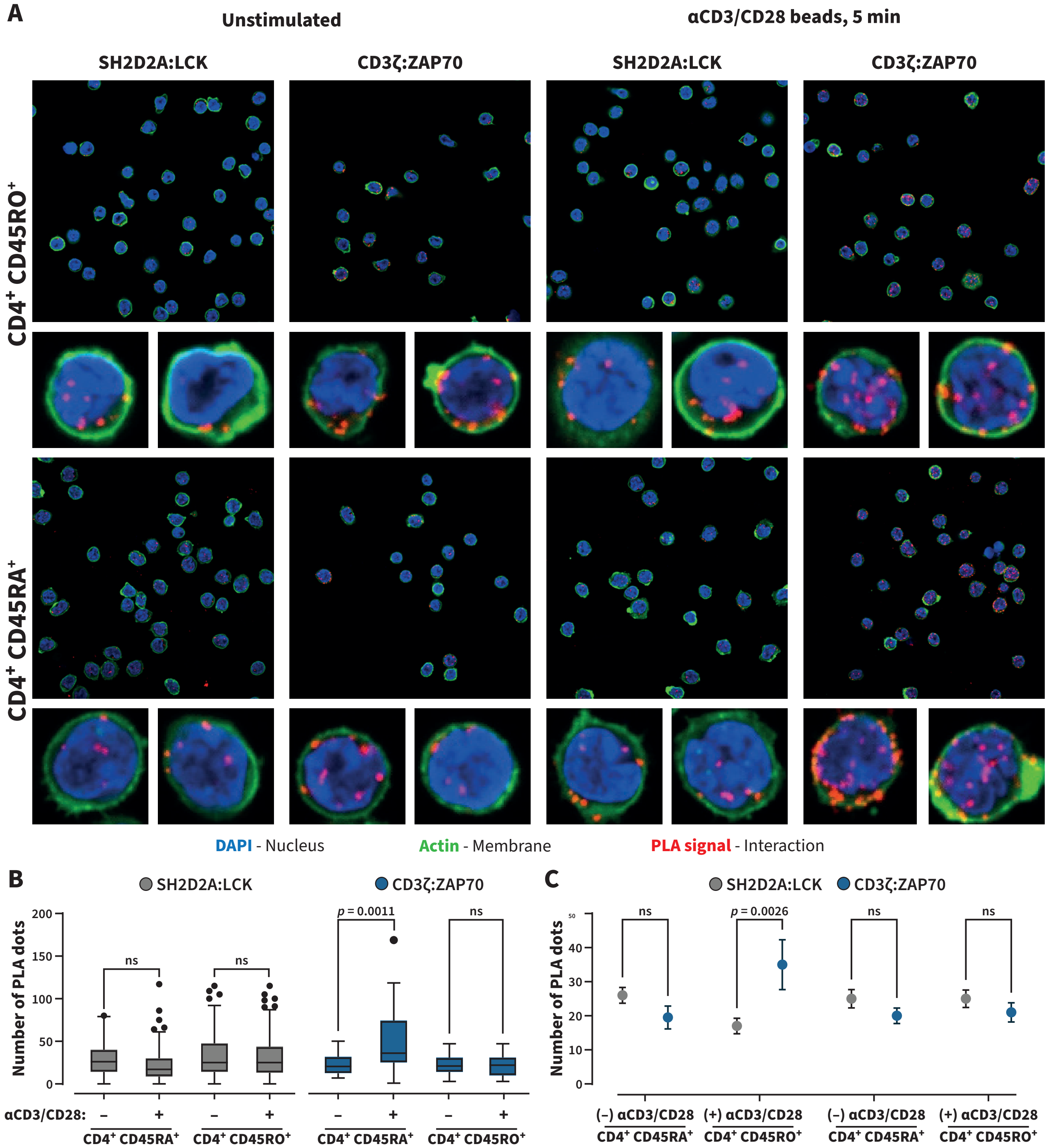


**Figure S1. The interaction of SH2D2A and LCK is not altered by αCD3/CD28 stimulation. (A)** Confocal images of proximity ligation assay (PLA) experiments performed on SH2D2A:LCK (1st and 3rd columns) and CD3ζ:ZAP70 (2nd and 4th columns) in memory CD45RO+ (top) or naïve CD45RA+ (bottom) CD4+ T cells, either stimulated for 5 min with αCD3/CD28 beads (right), or left unstimulated (left) – two representative single cells from the larger field images are shown below each image. **(B)** Box plots with Tukey whiskers comparing the number of PLA dots (and thus protein-protein interactions) per cell in each of the unstimulated vs. bead-stimulated samples in **(A)**, statistical significance was determined by analysis of variance (ANOVA) with Tukey corrections. **(C)** Summary plots of the average number of PLA dots per each sample, whiskers indicate the standard error of the mean (SEM), a two-way ANOVA with Šídák corrections was used to determine statistical significance.

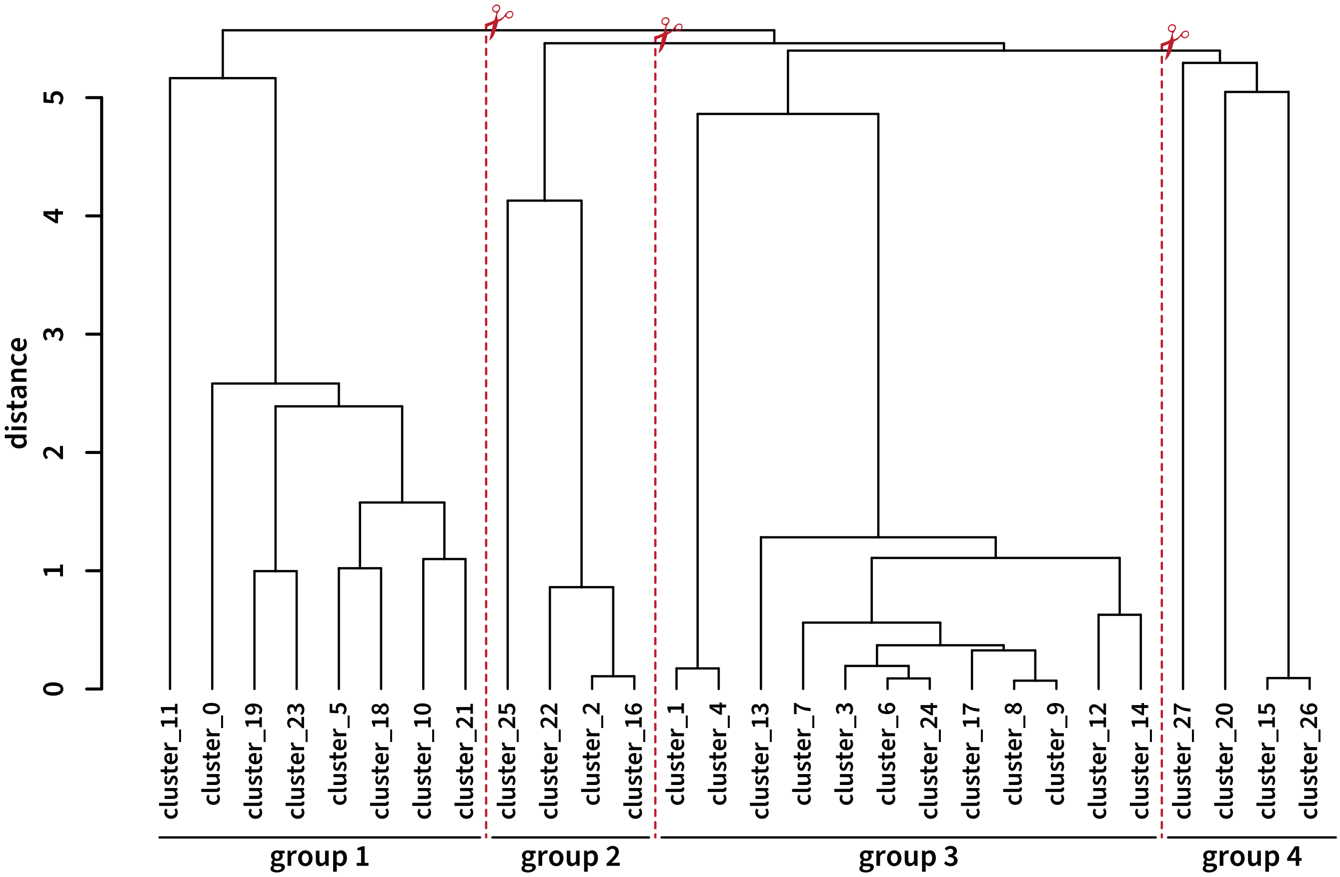


**Figure S2. Hierarchical cluster tree of the MAP-Liu dataset.** Tree depiction of distance matrix output of hierarchical cluster analysis of the dissimilarities between immune cell type content of the 27 clusters in **Figure 3A** as determined by the function *hclust()* from the library stats (v3.6.2). Higher branch points indicate a higher degree of dissimilarity between clusters. For **Figure 3B**, clusters were divided into four groups by cutting the three most-distant branches of the hierarchical clustering tree (red scissors).

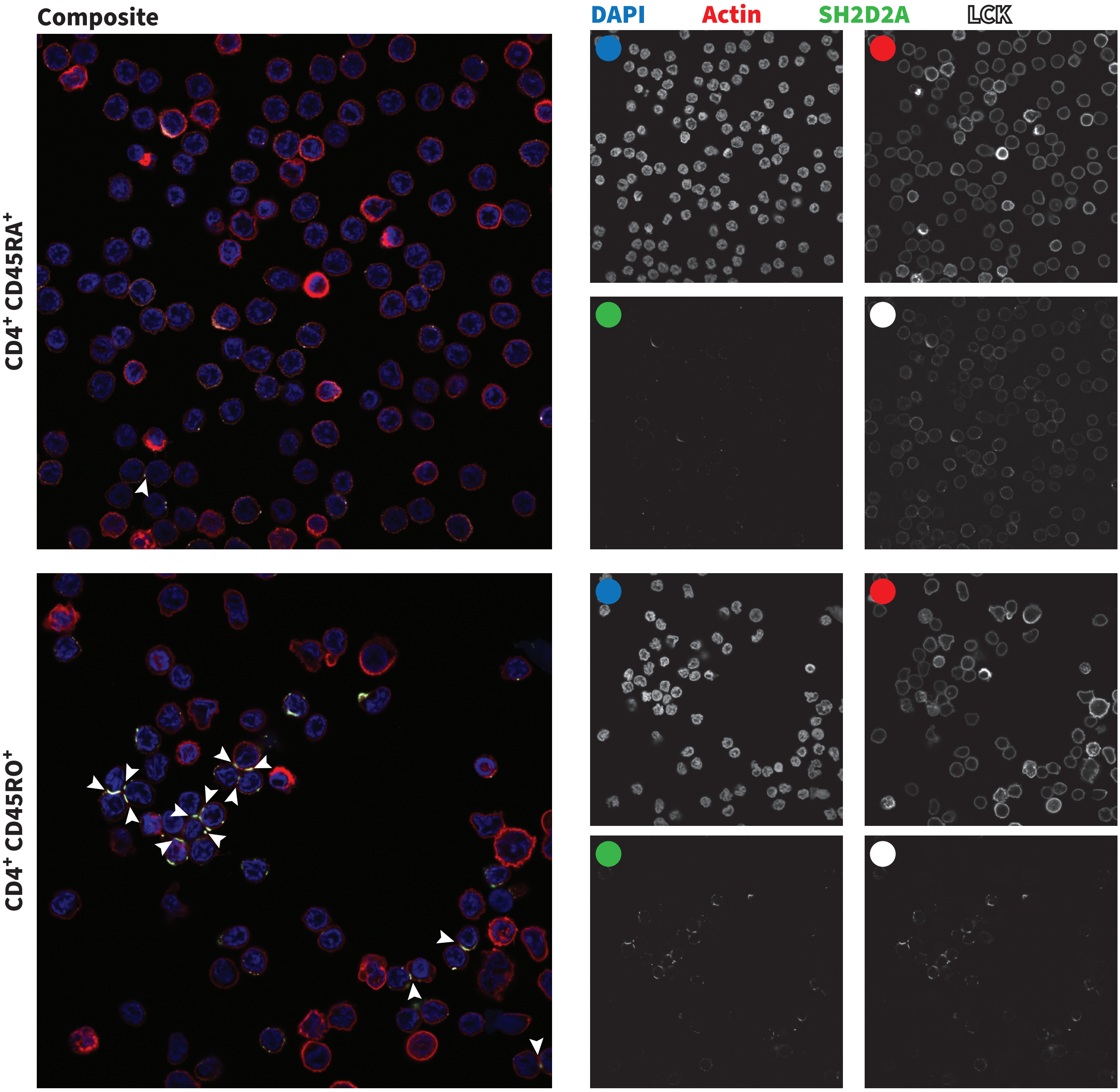


**Figure S3. CD4⁺ CD45RA⁺ cells display no clustering or synaptic accumulation of SH2D2A or LCK.** Confocal images of naïve CD45RA⁺ (top) and memory CD45RO⁺ (bottom) CD4⁺ T cells stained for DAPI (nucleus, blue), actin (membrane, red), SH2D2A (green) and LCK (gray). Images show the composite overlay of all four channels (left), as well as black-and-white images of the intensities of the four individual channels (right). Areas of abundance of LCK at T-T junctions are denoted with white arrows. Representative images from 1 donor (n = 5)

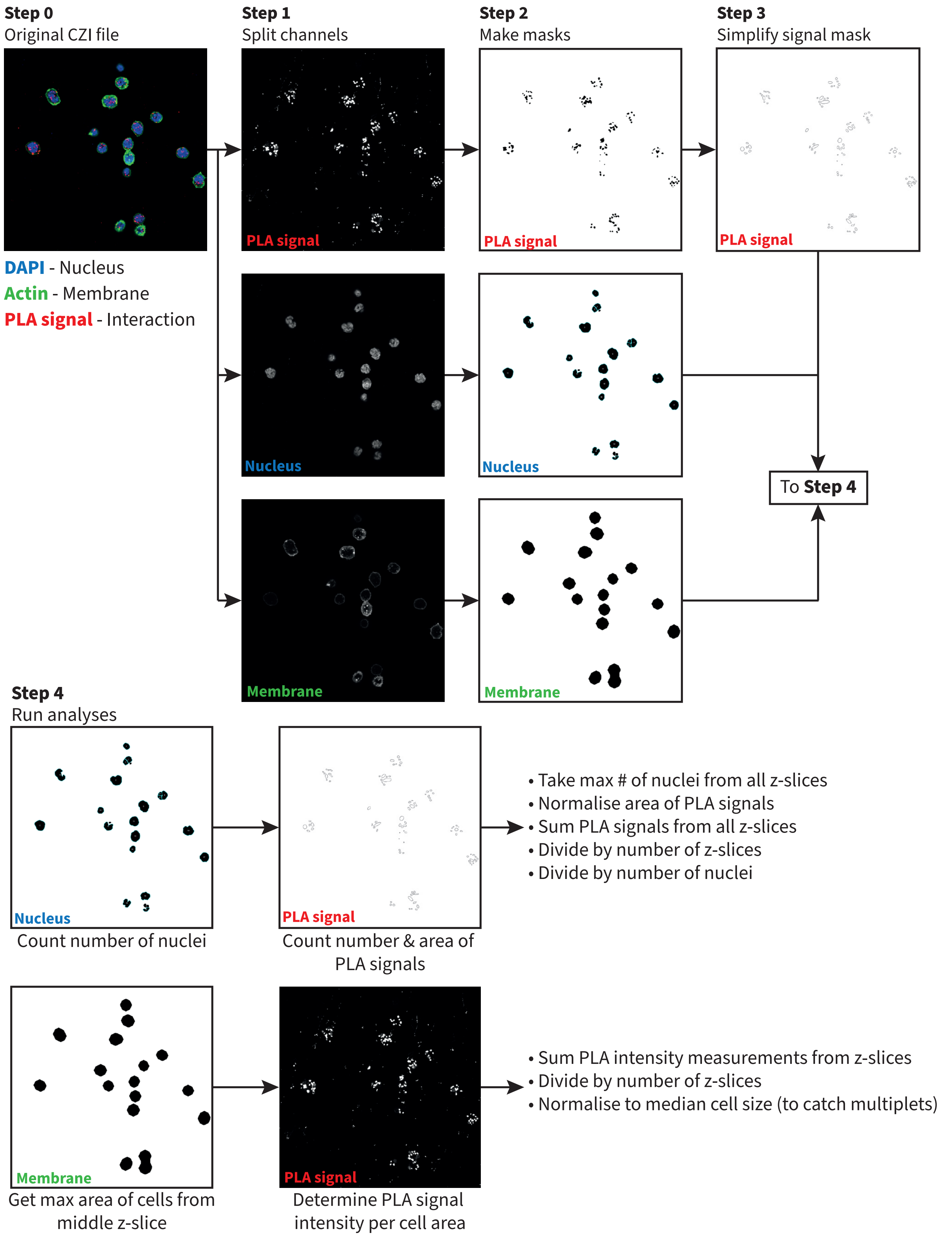


**Figure S4. Workflow depiction of the custom IJM script used to analyse confocal PLA images.** Depiction of the workflow carried out by the custom IJM script used to summarise confocal images taken following proximity ligation assay (PLA) experiments. Median Z-slice masks of the membrane channel were used to isolate single cells and carry out all analyses both on a global, image-wide scale, and on a per-cell basis.
